# Functionalized Calcium Carbonate Microparticles in Ethyl Cellulose Films: A Vehicle for Sustained Amoxicillin Release for Medical Applications

**DOI:** 10.1101/2025.02.19.639024

**Authors:** Petru Niga, Simone Sala, Jenny Rissler, Lina Nyström, Anna Fureby, Ulla Elofsson, Roger Roth, Patrick Gane, Joachim Schoelkopf

## Abstract

The continuous quest for materials capable of providing extended release of antimicrobial drugs is particularly important for indwelling medical applications. In this study, we utilized amoxicillin as a model active pharmaceutical ingredient (API) to investigate the feasibility of using porous media – specifically, functionalized calcium carbonate (FCC) microparticles – as a primary drug carrier embedded within an ethyl cellulose (EC) polymer film. Our main objective was to prolong and sustain the release of the API. The fabrication process of the microparticle containing film involved two key steps: loading the model API into the FCC particles and then embedding these loaded particles into the polymeric film. Amoxicillin was loaded into the FCC particles using a solvent evaporation method. Detailed characterization through Scanning Electron Microscopy (SEM), lab- and synchrotron-based XRD revealed that amoxicillin precipitated both inside and on the surface of the FCC particles, predominantly in an amorphous form. Additionally, ultraviolet-visible (UV-vis) spectroscopic data demonstrated an increased release rate from the particles compared to pure amoxicillin powder. Embedding amoxicillin pre-loaded porous FCC particles in the EC film led to a more rational sustained release compared with powder amoxicillin embedded directly in the film, advantageously delivering the same amount of amoxicillin over a longer period; a result that may be relevant for indwelling medical devices such as urinary catheters, vascular access devices or wound drains.

## 1. Introduction

Sustained drug release offers several advantages over conventional drug delivery systems. In conventional delivery, the blood plasma concentration of a drug typically increases abruptly after administration, peaking above the maximum desired level, which may lead to adverse effects. Subsequently, the concentration drops below the minimum effective level, rendering the drug ineffective, and increasing the chance to develop bacterial resistance (1). In contrast, sustained drug delivery acts to release the drug at a controlled rate over an extended period, enhancing therapeutic efficacy. This approach is particularly beneficial for patients with low compliance (2). Amoxicillin, a β-lactam penicillin derivative, discovered in 1958 by Doyle, Nayler and Smith (3), that can withstand the acidic environment of the stomach, is widely used to treat various bacterial infections and is one of the most commonly prescribed antibiotics in primary care settings (4). However, the extensive use of antibiotics has led to the rise of antimicrobial resistance, which is one of the most pressing challenges in modern medicine (2). To combat this, researchers are focusing on discovering new antibiotics and developing formulations that limit antimicrobial resistance. Sustained drug release systems are of particular interest because they can maintain drug levels above the minimum inhibitory concentration, without increasing the total amount of drug released, thereby reducing the likelihood of resistance development.

Sustained release formulations of amoxicillin for oral delivery have been extensively studied. For instance, the mucoadhesive sustained release of amoxicillin from Eudragit RS100 microspheres demonstrated that the release profile depended on both the mucosa and the swelling ability of the polymer, sustaining release for approximately 12 h (5). Additionally, a 24-h sustained release was achieved by loading amoxicillin into composite hydrogels made of poly(acrylamide) and starch, where release kinetics were sensitive to both temperature and pH (6). Extended release has also been achieved by coating amoxicillin particles with ethyl cellulose (EC), with additional chitosan or chitosan-cyclodextrin coatings further prolonging release (7).

For indwelling applications, sustained release must be controlled over longer periods to maintain therapeutic efficacy. For example, hydroxyapatite nanoparticles loaded with amoxicillin and coated with polyvinyl alcohol or sodium alginate achieved a sustained release over 30 days, which was explored for treating bone infections (8).

Currently, significant efforts are directed at developing new indwelling urinary catheters (IUCs) designed to prevent bacterial infections, a major healthcare and patient well-being concern. Despite advances in antifouling and antibacterial catheter coatings, translating these concepts into clinical practice remains challenging due to discrepancies in experimental evaluations. For a comprehensive review of materials and their effectiveness, readers are referred to Andersen and Flores-Mireles, 2020 (9).

In this study, we explore the feasibility of using a polymeric matrix containing porous particles loaded with amoxicillin as a model antibiotic to form a platform to exemplify the sustained release of an active pharmaceutical ingredient (API), intended, for example, for use in catheter applications. This application would strongly benefit from extended-release time, which cannot be achieved by simply mixing the active ingredient into the polymer matrix (also confirmed in this study). The porous particles used in this study are functionalized calcium carbonate (FCC). They are composed of calcium carbonate surrounded by an intertwined layer of calcium phosphate platelets, offering enhanced specific surface area and high interconnected porosity (10–12).

In the first step of this study, we investigated the loading and release profile of amoxicillin from the FCC particles. The state of the precipitated amoxicillin in the FCC was analyzed using SEM and lab- and synchrotron-based XRD. In the second step, the amoxicillin-loaded FCC particles were uniformly dispersed in ethyl cellulose (EC) and cast into a film. Additionally, the release profile of amoxicillin from this film in water was evaluated using UV-vis spectrometry.

## 2. Materials and Methods

### 2.1. Materials

Amoxicillin was sourced from Sigma Aldrich with a purity higher than 99.9 %. Ethanol, methanol, and methyl ethyl ketone (MEK) were also sourced from Sigma Aldrich. Porous functionalized calcium carbonate (FCC) particles were supplied by Omya International AG, Switzerland. In the loading and release experiments we used the 0–180 μm (here referred to as FCC fines) and 180–710 μm (here referred to as FCC granules) fractions of the laboratory listed TP_500DC-BER (100 % roller compacted FCC batch TP2907/J1) sample. These two fractions are derived from Omyapharm^®^ FCC 500-OG (here referred to as Omyapharm), which was used in the film manufacturing experiments. This starting material has the weight median particle size *d*_50 w/w%_ of 6.6 μm, specific surface area BET of 53 m^2^/g and apparent bulk density of 0.13 g/cm^3^. Particles were dried at 200 °C for 2 h before processing. Ethyl Cellulose N100 (Ethoxyl grade 48–49.5 %) was generously provided by IMCD Group. Water was sourced from a MilliPore RiOs-8 system. Reference amorphous amoxicillin used in synchrotron experiments was prepared by solvent evaporation method in a rotary evaporator.

### 2.2. Particle Loading

To load amoxicillin into the porous FCC particles, the amoxicillin must first be dissolved in a solvent with a low boiling point. This solution is then mixed with the FCC particles, and the solvent is allowed to evaporate. During evaporation, amoxicillin precipitates both within the pores and on the surface of the FCC particles. In this study, a 3 g/L solution of amoxicillin in an ethanol/acetone mixture was prepared. This solution was then added to a predefined amount of particles (for 30 % and 15 % loading – 7 g FCC and 8.5 g FCC, respectively, per 1 L solution) and the solvent was evaporated using a rotary evaporator (30 °C and 400-150 mbar).

### 2.3. Material characterization and techniques

#### 2.3.1. Loading Level and Release Studies

The loading level and extent of amoxicillin release in water were determined using a thermogravimetric analyzer (TGA 2 Stare System, Mettler Toledo, Columbus, OH, USA) with a STARe System software. The system was heated from 25-800 °C at a rate of 20 °C/min. The amount of amoxicillin in the FCC particles was measured by analyzing the material’s weight loss as it was heated.

#### 2.3.2 Imaging and Spectroscopic Analysis

Imaging was performed using a Quanta™ FEG 250 Scanning Electron Microscopy SEM (FEI Instruments, Hillsboro, OR, USA). The concentration of amoxicillin in water was measured using a Lambda™ 650 UV-vis spectrometer (PerkinElmer, Shelton, CT, USA) by recording absorbance at 272 nm, with concentration determined by interpolation from a calibration curve.

#### 2.3.3. Crystal Structure

To investigate the crystal structure of amoxicillin, two X-ray diffraction (XRD) instruments were employed:

i. Lab Bench X-Ray Diffractometer: An X’Pert PRO diffractometer (Malvern Panalytical, Malvern, UK) was used under the following conditions: temperature at 26.5 °C, a 1 mm-thick compacted powder sample, Bragg reflection geometry, spinning sample holder, and a zero-background holder with a Si wafer. Use of the Cu anode’s Kα emission line resulted in an incident photon energy of 8.05 keV.
ii. Synchrotron Hard X-Ray Nanoprobe Beamline: Measurements were conducted at the NanoMAX beamline at MAX IV, Sweden (13, 14). The incident photon energy was set to 15 keV and the XRD signal was collected using the Pilatus3 X 1M detector (Dectris, Baden, Switzerland), with 981 × 1043 pixels, each measuring 172 μm. To achieve a higher photon flux (1.1 × 10¹⁰ photons/s), the beamline’s slits were slightly opened, resulting in a partial loss of coherence and a focused X-ray beam of 80 nm × 80 nm. Two-dimensional (2D) XRD measurements were carried out by scanning the sample through the nano-focused X-ray beam with a step size of 80 nm, matching the focal spot size. This

generated 2D maps with a lateral resolution of 80 nm and for which a diffraction pattern is available at every pixel. Simultaneous detection of the characteristic X-ray fluorescence (XRF) emission from sulfur was carried out using the beamline’s single-element silicon drift detector (SDD) (RaySpec, High Wycombe, UK). As amoxicillin was the only compound containing sulfur within the investigated samples, sulfur distribution was used as proxy for amoxicillin distribution. For the experiments, single particles were deposited by electrostatic precipitation from air on thin Si_3_N_4_ membranes (1 µm thick).

### 2.4. Film preparation and fabrication

Film preparation was carried out using organic solvent evaporation to avoid high-temperature curing, which could degrade the thermally sensitive amoxicillin (15). To minimize the dissolution of the active ingredient into the film matrix, an organic solvent mix (toluene: ethanol) was selected that dissolves ethyl cellulose (EC) efficiently while offering minimal solubility for amoxicillin.

For film production, Omyapharm particles (the starting material for FCC fines and granules) were loaded with 30 w/w% amoxicillin using a solvent evaporation process with a MEK/methanol mixture. For the polymer solution, 5 g of EC-N100 grade polymer was dissolved in 100 mL of an 80:20 w/w% toluene:ethanol mixture, resulting in a clear, viscous solution. Upon adding the amoxicillin-loaded Omyapharm particles, a white dispersion was obtained, as illustrated in Figure 1A. This suspension did not show any signs of agglomeration or sedimentation for more than 1 h, which was sufficient to produce the films. The films were applied onto a low-density polyethylene (LDPE) substrate using a hand coater (Erichsen – Germany) with a 200 μm indenture. Note that the final dried film thickness was not measured; throughout this paper, we refer to film thickness in relation to the size of the edge indenture (e.g., 200 μm).

**Figure 1.**
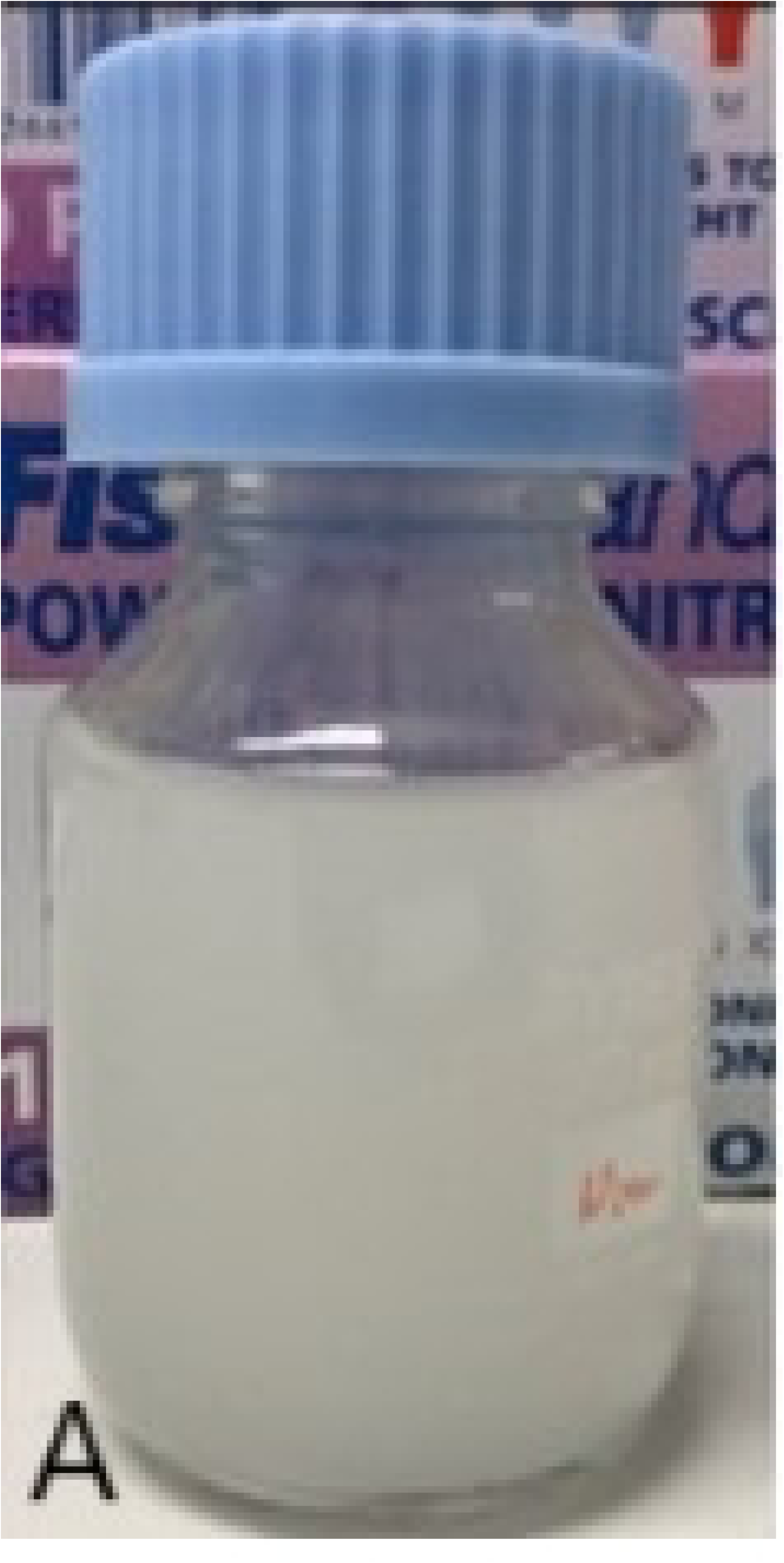
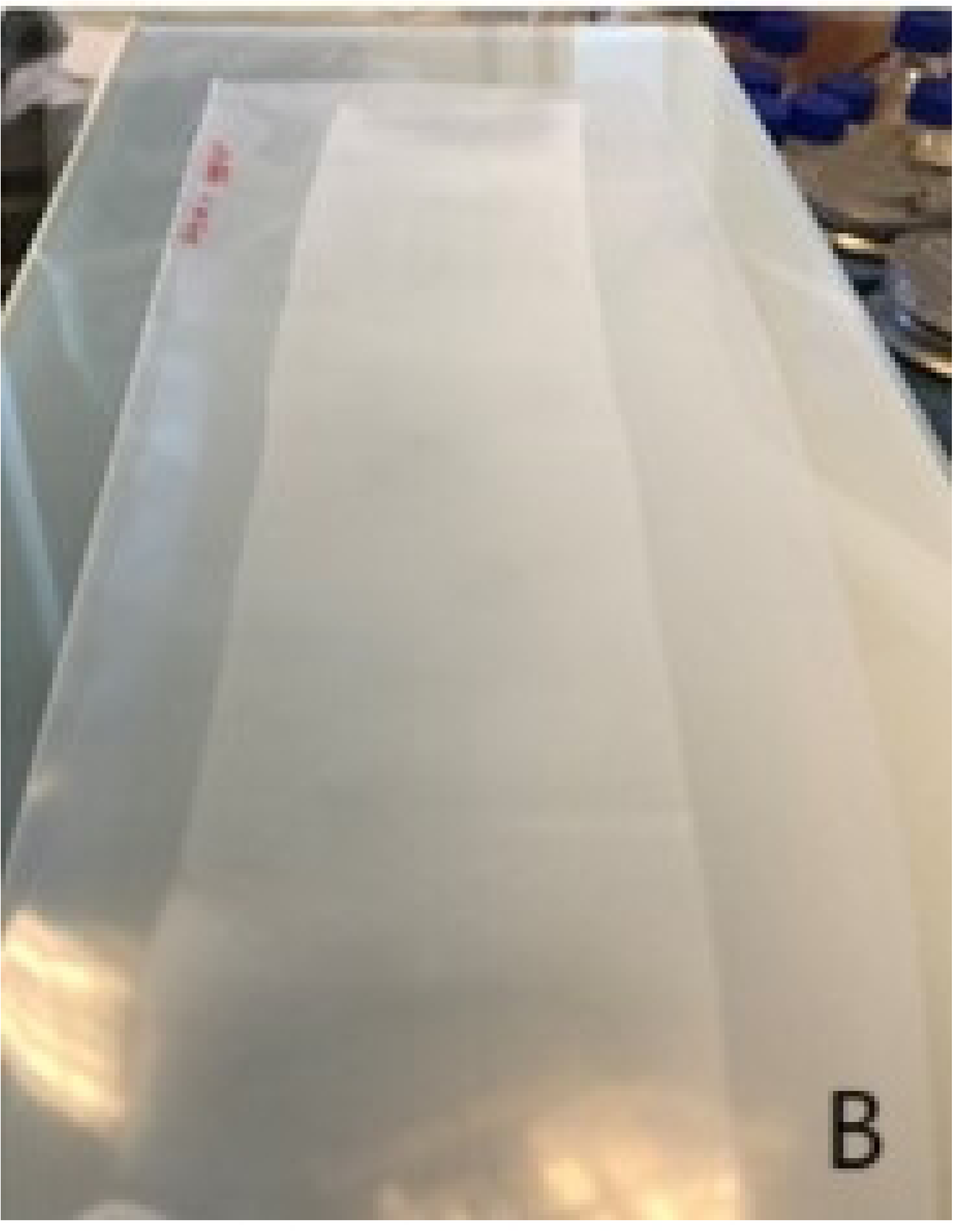
A) 30 w/w% amoxicillin-loaded Omyapharm suspended in a solution of EC-N100 dissolved in toluene: ethanol mixture, and B) a film coated on a LDPE substrate.

The maximum particle content (Omyapharm) within the total solid content used in the dispersion formulation was defined as the value at which the resulting film adhered well to the substrate over time without peeling. This maximum was found to be 10 w/w% of total solid content for all film thicknesses tested. Additionally, it was visually observed that the distribution of Omyapharm particles throughout the film was uniform at this concentration. Figure 1 shows 30 w/w% amoxicillin-loaded Omyapharm particles suspended in a solution of EC-N100 (1A), and the final film coated on the LDPE substrate (1B).

### 2.5. Experimental setup for release studies

All release experiments were carried out in Milli-Q water at a starting pH of 6.99 and a temperature of 22 °C. For the release from particles, the weight of the samples was chosen to maintain sink conditions throughout the duration of the experiments. Particles and pure amoxicillin were placed in a beaker, containing 75 mL of Milli-Q water and stirred at 100 min^−1^(RPM). The dissolution media was circulated via a peristaltic pump into the UV-spectrometer and returned back into the beaker. Absorbance was measured at 272 nm and concentration was calculated using a linear calibration curve.

For the experiments testing release from the film, 200 μm thick films containing 10 w/w% of particles loaded with amoxicillin were used. The measurements were carried out in 5 mL vials and samples (1 mL) were manually withdrawn at specified time intervals and returned after UV-vis measurement. The film samples were not agitated.

To adjust the surface area exposed to water, a special configuration of the film was employed, such that the film was perforated with a pin and a wire passed through the holes to keep the adjacent film roll apart and allow water to penetrate, as seen in Figure 2. The film surface area exposed to water in this way was calculated to be 5.4 cm^2^/mL.

**Figure 2.**
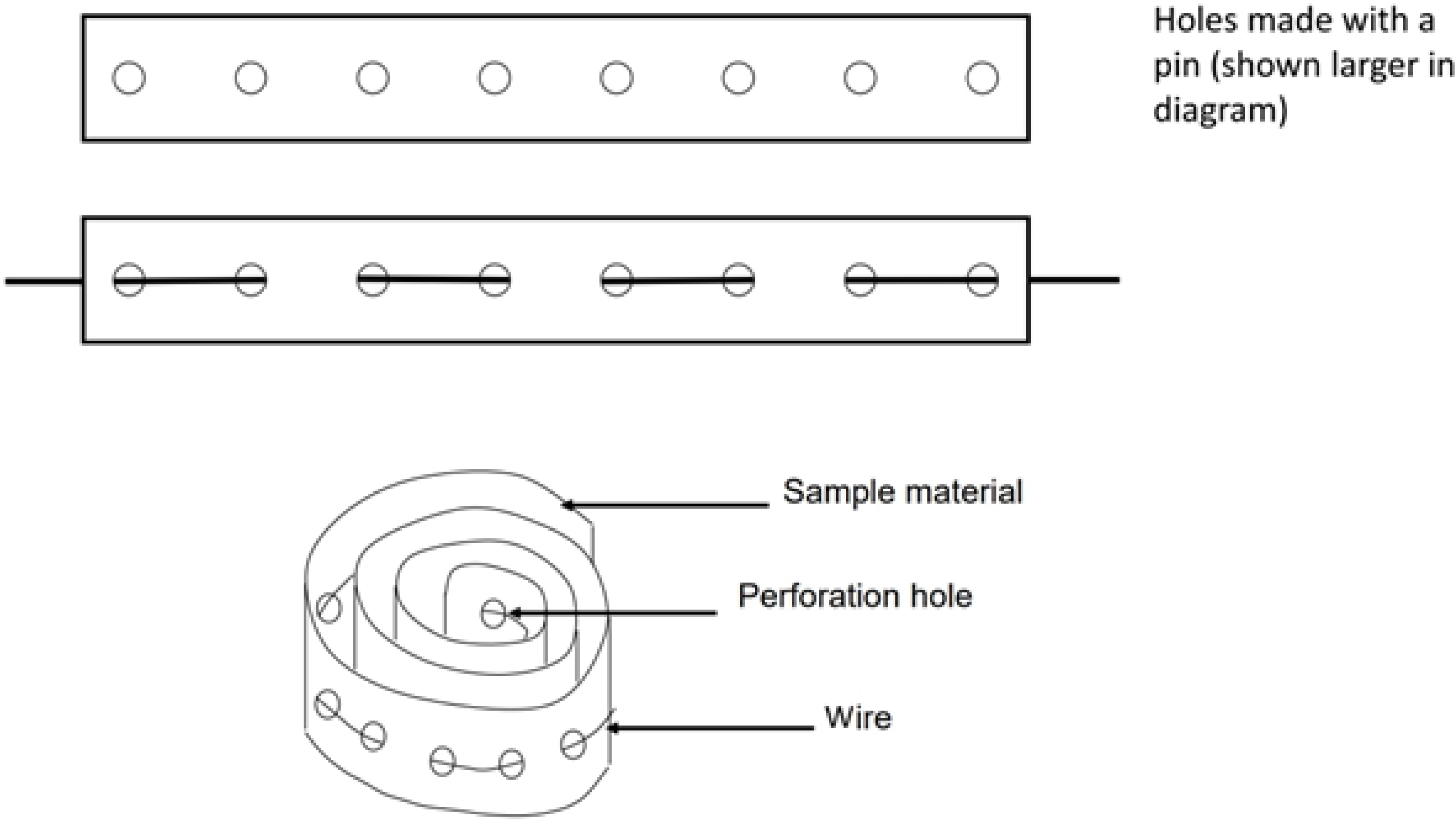
Configuration of the film placed in the release beaker.

## 3. Results and discussion

### 3.1. Drug-load in porous particles

Amoxicillin, the model API, was loaded into the porous particles using the solvent evaporation method. Loading levels were quantified via TGA, as shown in Table 1. The measured loading levels were slightly lower than the calculated values, likely due to amoxicillin precipitating on the walls of the rotary evaporator during solvent evaporation.

**Table 1.** Calculated and measured amoxicillin loading values in porous particles by TGA.

| Method | Amoxicillin concentration in |  |  |  |
| --- | --- | --- | --- | --- |
|  | FCC fines / w/w% |  | FCC granules / w/w% |  |
| Calculated | 15 | 30 | 15 | 30 |
| Measured | 13 | 22.5 | 12 | 27 |

As shown in Figure 3, the release profile exhibited a two-stage pattern: an initial rapid release within the first 2-3 min, followed by a slower release over the next 30 min, approaching (but not reaching) a plateau. After 30 min, the 30 w/w% and 15 w/w% amoxicillin-loaded FCC fines reached concentrations of 99 w/w% and 94 w/w% of the total amount, respectively. After 24 h, all of the loaded amoxicillin was released into the solution, as confirmed by TGA (see Supplementary Information). The release behavior from FCC granules followed a similar trend (also recorded in the Supplementary Information). Even though in both cases the amoxicillin was fully released in water, the release curves of the two samples show different trends. That may be because the amoxicillin on the outer part of the particles is released more easily (which is assumed to be more in the 30 w/w% case) while amoxicillin adsorbed in the inner part, close to the pore wall surface, is released more slowly.

**Figure 3.**
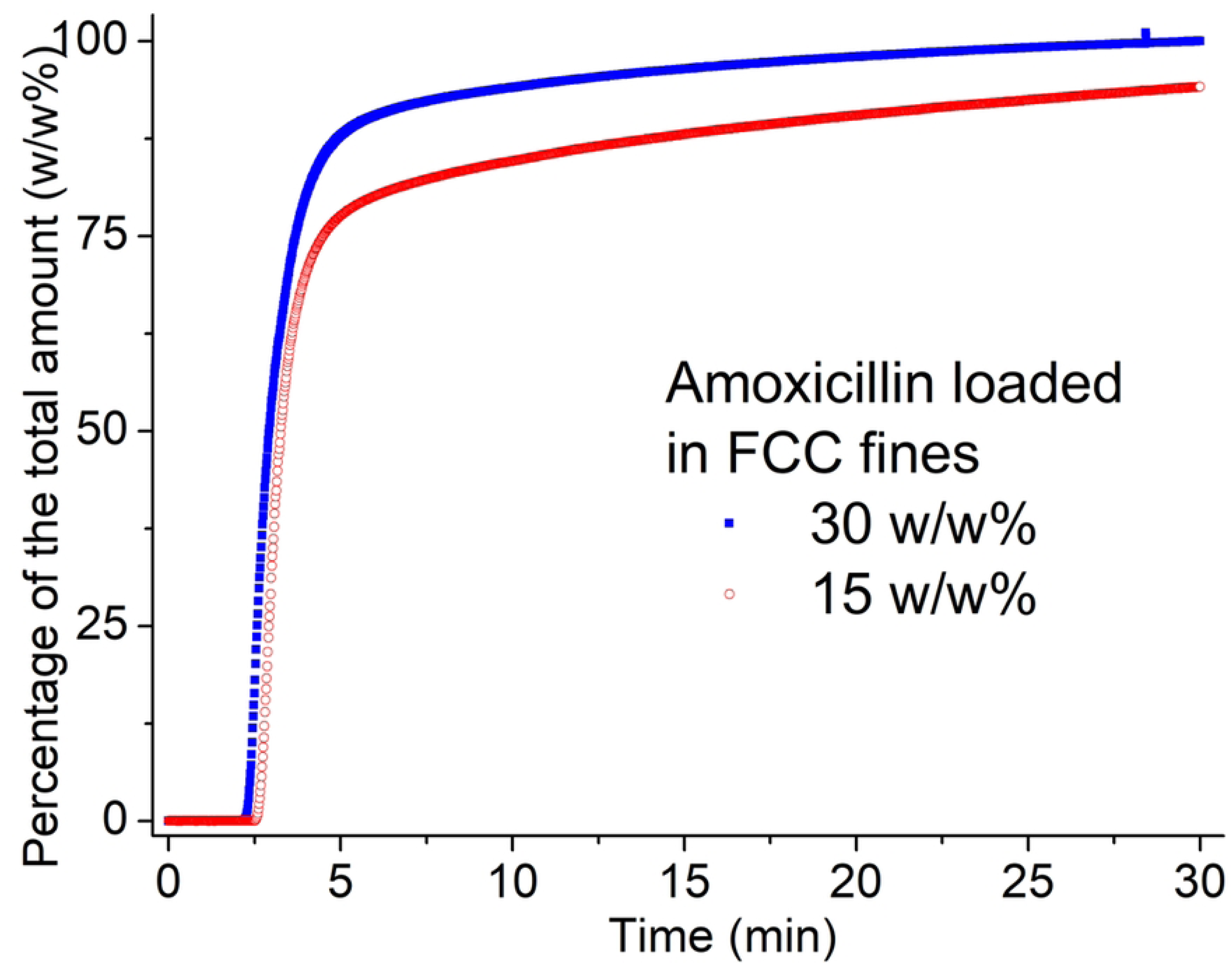
The amoxicillin release curves in water from 15 w/w% (red circles) and 30 w/w% (blue squares) amoxicillin loaded FCC fines.

To assess the benefits of loading amoxicillin into porous media (specifically FCC granules), a comparative experiment was conducted. In this experiment, the release profile of amoxicillin-loaded FCC granules was compared to that of amoxicillin powder (as purchased) under identical conditions. For both forms, the quantity of amoxicillin added to the beaker was calculated to be below the solubility limit (1 mg/mL) if fully released. As illustrated in Figure 4, during the initial rapid release phase (from 3 to 3.6 min), the slope of the fitted release curve was slightly higher for the amoxicillin released from FCC granules (1.22 w/w%/min) compared to the powder form (1.083 w/w%/min). Additionally, while all the amoxicillin loaded in FCC granules was released within the experiment’s duration, only 95 w/w% of the as-purchased powder form dissolved. These findings indicate that amoxicillin loaded in FCC granules not only releases more quickly but also more completely in water compared to the powder form. This suggests an enhanced dissolution rate, which is not typically observed for crystalline amoxicillin.

**Figure 4.**
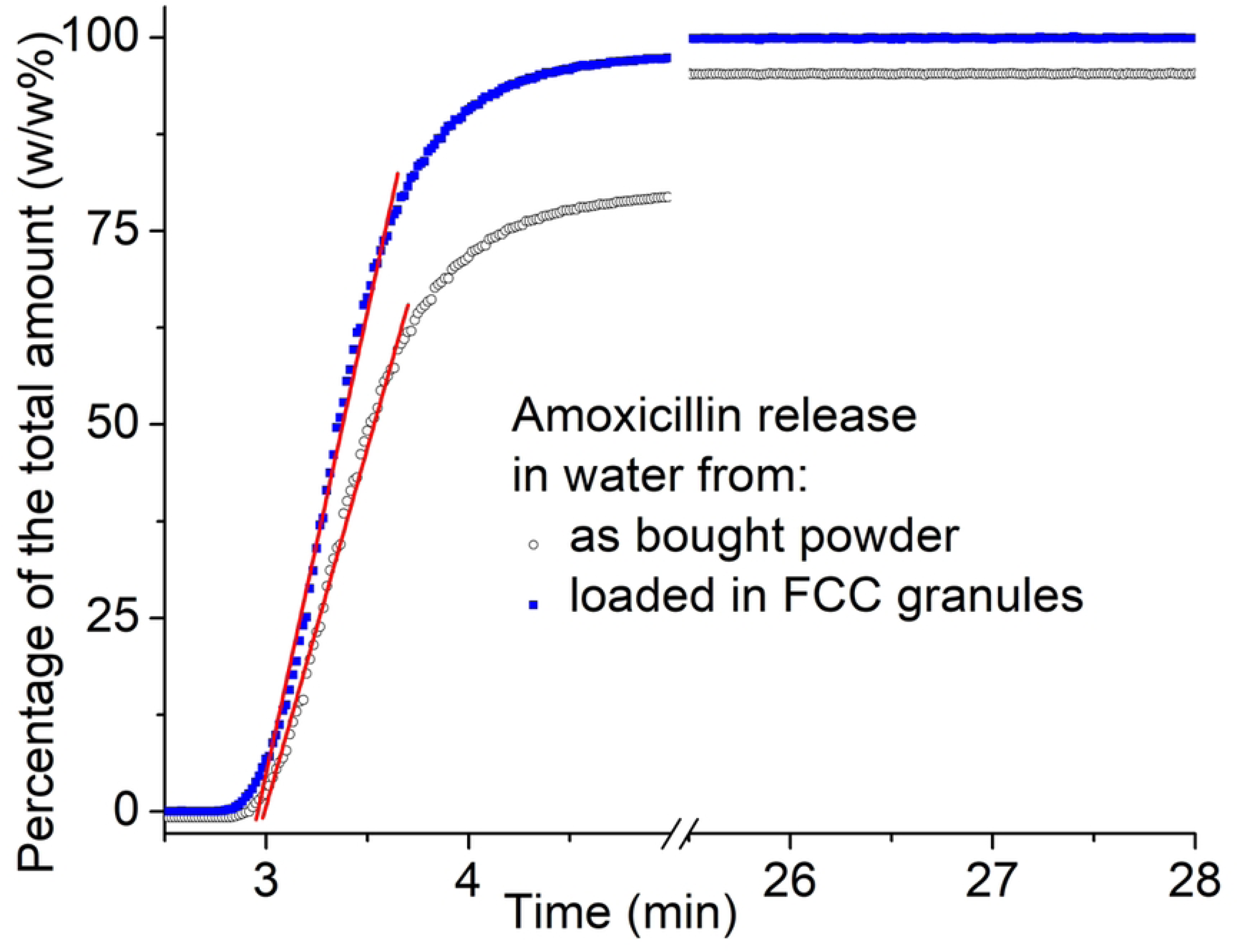
The estimated slope of the amoxicillin release in water from FCC granules (blue squares) compared to as bought powder (black circles). The red line is the fitting curve from 3 to 3.6 min.

### 3.2. XRD experiments

To understand the underlying mechanisms influencing amoxicillin release, the structural characteristics of the loaded amoxicillin were examined using both lab bench and synchrotron-based XRD (2D-mapping).

#### 3.2.1. Lab bench XRD experiments

For the XRD analysis, three samples were investigated: (i) amoxicillin powder, (ii) amoxicillin loaded into FCC fines, and (iii) FCC fines as a reference. The crystallite cell parameters of the amoxicillin powder were determined as follows: unit cell volume = 1781.47 Å³, with parameters a = 14.76 Å, b = 20.85 Å, c = 5.79 Å, and β = 92.53°. These values closely align with the orthorhombic single crystals reported in the literature (16–18).

The diffraction pattern of amoxicillin powder, shown in Figure 5, exhibits numerous peaks in the *q* = 7 – 46 nm^−1^ range, indicating a highly crystalline phase. The recorded intensities, reaching up to 400k counts, exceed those reported in previous studies due to the extended data collection time employed in this work (7). In contrast, the diffraction patterns of the FCC fines reference material and amoxicillin-loaded FCC fines display only 15 to 20 significant peaks in this region, suggesting a lower degree of crystallinity.

**Figure 5.**
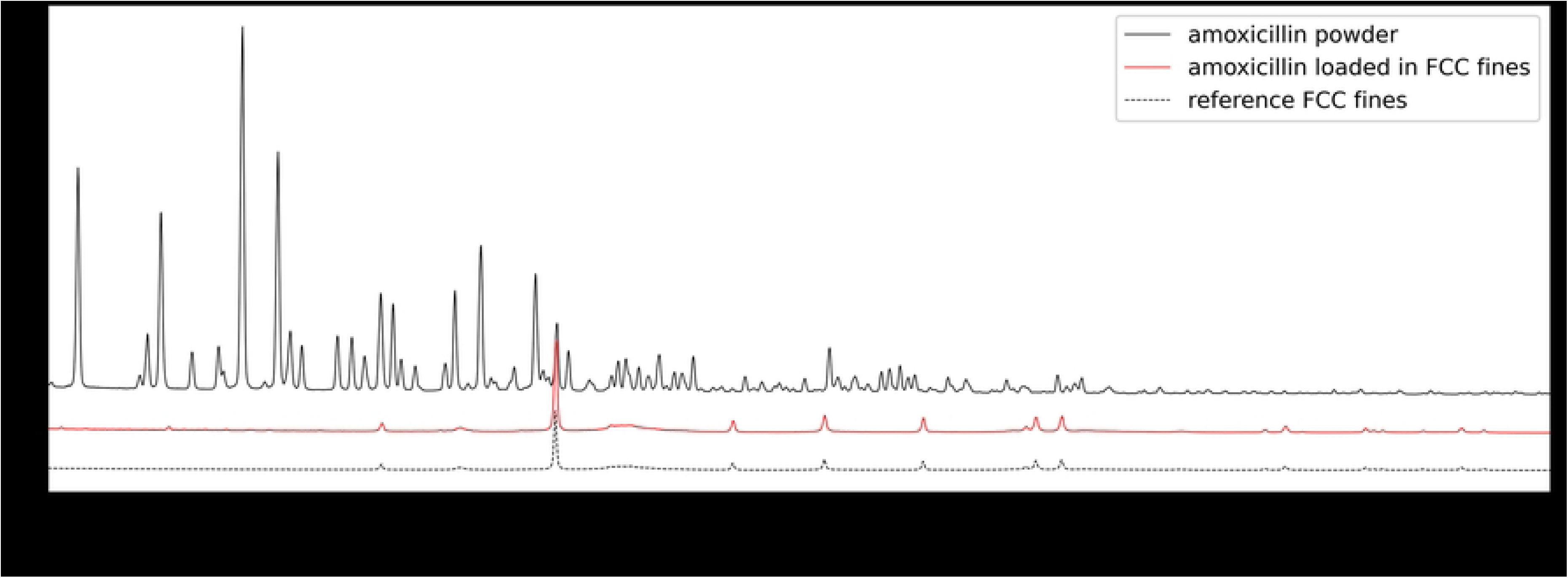
Diffraction pattern for amoxicillin loaded in FCC fines (continuous red curve), amoxicillin powder (continuous black curve) and FCC fines reference (dashed black curve). Curves have been offset by a constant value for easier visualization.

In fact, most of the features above 15 nm^−1^ are associated with the FCC fines reference material, while the features below 15 nm^−1^, about 5-6 minor peaks, are associated with the loaded amoxicillin. This indicates that the amoxicillin loaded onto the particles has very limited long range crystalline order (19). Generally, amorphous materials have no peaks in diffraction patterns. However, it must be remembered that the material mass and electron density of amoxicillin in comparison to FCC is on average considerably less. The intensity of the visible features may, nonetheless, help elucidate whether the loaded amoxicillin is amorphous or manifesting an alternative structure, such a nanocrystals or a liquid crystal mesophase. Both amorphous and nanocrystalline phases contain no long-range order, meaning that there are no coherent multiple regular crystalline planes to diffract X-rays. For amorphous materials the incident X-rays are scattered isotropically and there are no sharp peaks in the diffraction pattern. In other words, long range ordered crystalline parts give sharp narrow diffraction peaks and a truly amorphous component would give a broad background contribution (19), which is usually termed a “halo”. The lack of sharp crystalline peaks or a significant amorphous halo cannot be attributed to an exceedingly weak interaction of X-rays with the samples. Thus, it might be more appropriate to consider the lack of an amorphous background in the spectrum in this case as an indication rather of nanocrystallites (lacking longer range order, with exceptionally high surface area) or a mesophase state exhibiting order in only one (nematic) or maximally two dimensions (smectic). These more complex structures could account for both the lack of diffraction peaks and the absence of amorphous background together with a high rate of solubility related to either the extremely high nanocrystalline or mesophase surface area.

The significant loss of crystallinity is evidenced by the 25-fold reduction in peak intensity when amoxicillin is loaded into the pores and onto the surfaces of FCC fines. Given that the total amount of the amoxicillin exposed in the two experiments was the same (pure amoxicillin and amoxicillin loaded into the FCC fines) and the fact that the signal intensity is proportional to the amount of crystalline material, a 30 w/w% decrease in material should result in only a one-third reduction in signal intensity, not a 25-fold decrease (20, 21). This stark reduction indicates a substantial loss of long-range crystallinity. Similar reductions in crystallinity were observed in other samples, including FCC fines loaded with 15 w/w% and 30 w/w% amoxicillin and FCC granules loaded with 15 w/w% amoxicillin, respectively (data available in the Supplementary Information).

Given that the XRD diffractogram of the loaded amoxicillin did not display the characteristic amorphous “halo” but a significant reduction in peak intensity compared to pure amoxicillin, the structural state of the amoxicillin within the porous material could, therefore, not be conclusively determined. To investigate further whether the loaded amoxicillin in the confined space of the porous FCC material is amorphous, nanocrystalline or mesophase, we conducted experiments with higher sensitivity and spatial resolution using the hard X-ray nanoprobe beamline, NanoMAX, at MAX IV in Lund.

#### 3.2.2. Synchrotron-based XRD experiment

For the synchrotron-based XRD experiments, four different samples were analyzed: (i) amoxicillin powder, (ii) amoxicillin loaded into FCC fines (30 w/w%), (iii) unloaded FCC fines, and (iv) reference amoxicillin precipitated by solvent evaporation. As expected, the diffractogram of the pure amoxicillin powder (continuous black line in Figure 6 A) exhibits intense peaks, characteristic of its crystalline structure. Conversely, the amoxicillin precipitated by solvent evaporation (dashed red line in Figure 6 A displays no discernible peaks, confirming a long-range disordered state. To facilitate comparison, each diffractogram was rescaled by dividing it by its average intensity within the low *q* range (9-13 nm^−1^) preceding the region featuring diffraction peaks. This normalization enables a clearer comparison of the structural features between the different samples.

**Figure 6.**
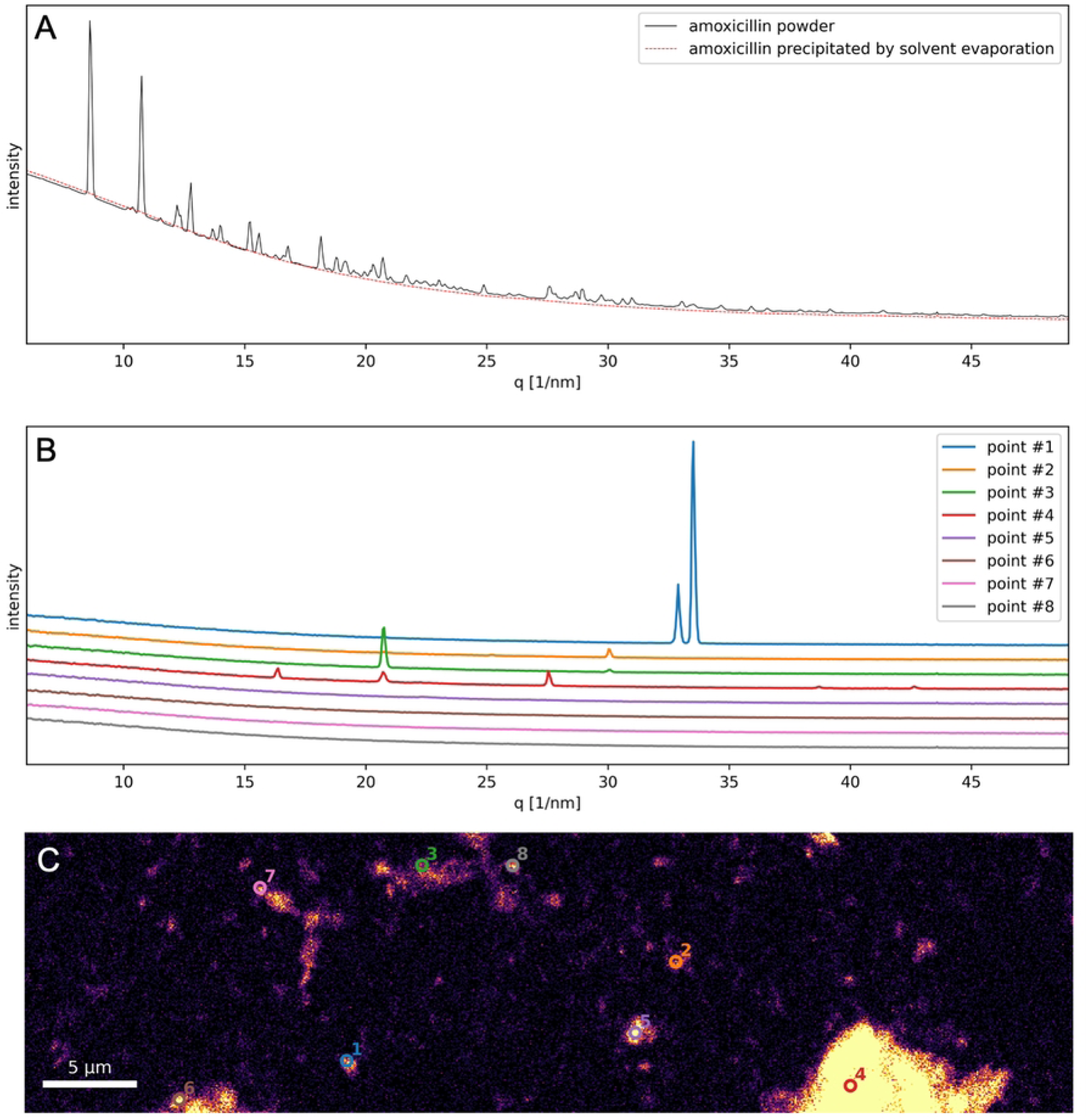
Normalized diffractograms of pure amoxicillin as compared to amoxicillin precipitated by solvent evaporation (A) and loaded in FCC fines (B); diffractograms in (B) have been offset by a constant value for easier visualization. High resolution map of the amoxicillin loaded in FCC fines (C): bright pixels indicate the presence of amoxicillin. The diffractograms displayed in (B) are extracted from the points annotated in (C).

The diffractograms from amoxicillin loaded in FCC fines exhibited significantly lower intensity features and only for a few scanning positions (see points 1-4 in Figure 6 B-C). This suggests a predominant loss of crystallinity in the amoxicillin loaded into the particles. The bright regions from Figure 6 C correspond to S-rich regions and therefore indicate the presence of amoxicillin: an inhomogeneous distribution with clusters of variable size is apparent. Interestingly, most pixels revealing a significant presence of amoxicillin – of which only 8 selected examples are displayed in Figure 6 B – correspond to diffractograms without visible diffraction peaks thus indicating that the majority of the amoxicillin present in this sample is in an amorphous state.

#### 3.3. Scanning Electron Microscopy

The SEM images presented in Figure 7 provide further support for the findings from the amoxicillin release studies and the X-ray experiments. The pure amoxicillin exhibits elongated, rod-like crystalline formations, which resemble elongated prismatic shapes. In contrast, the amoxicillin precipitated within the pores or on the surface of the FCC fines displays a distinct snowflake-like structure.

**Figure 7.**
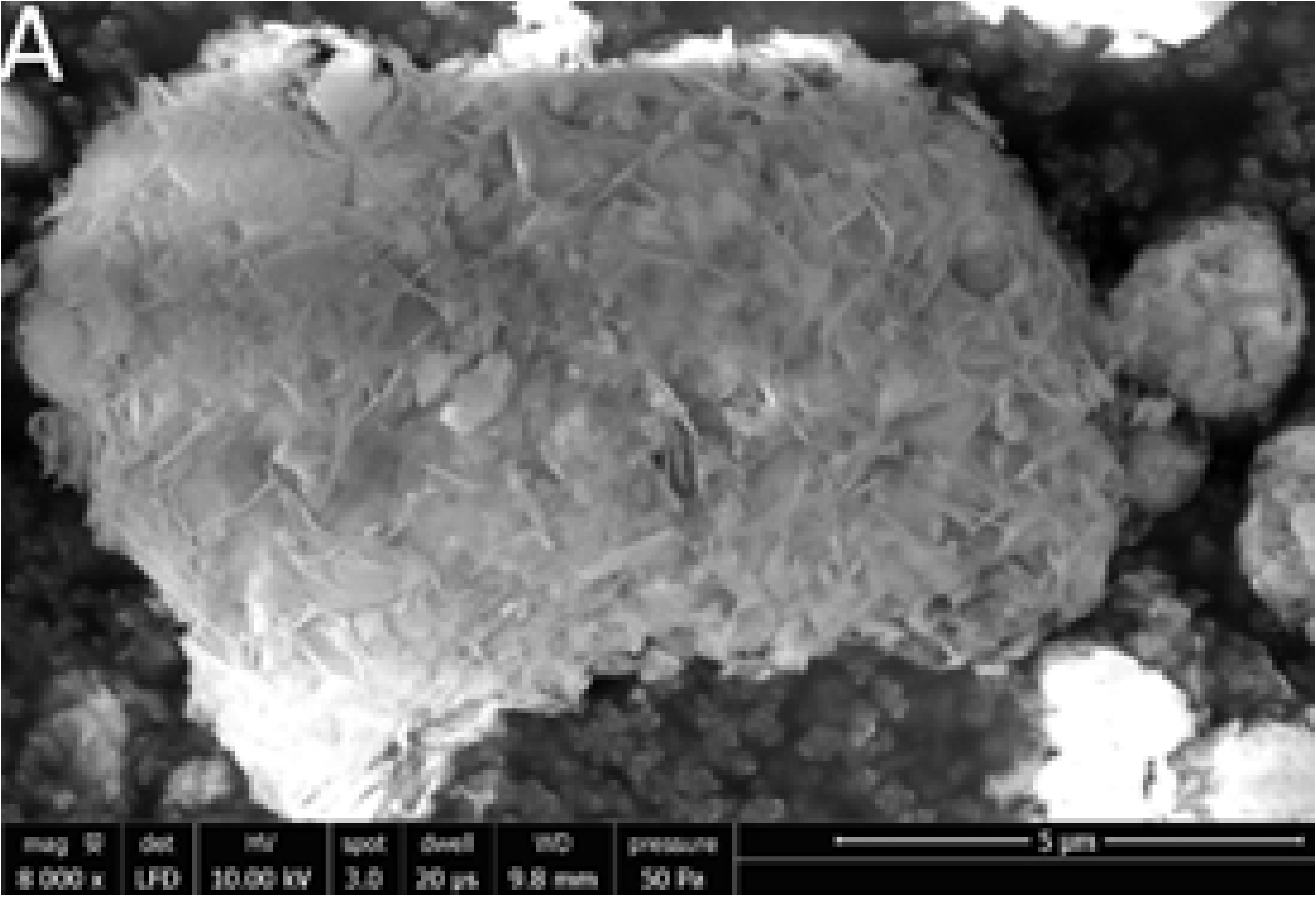
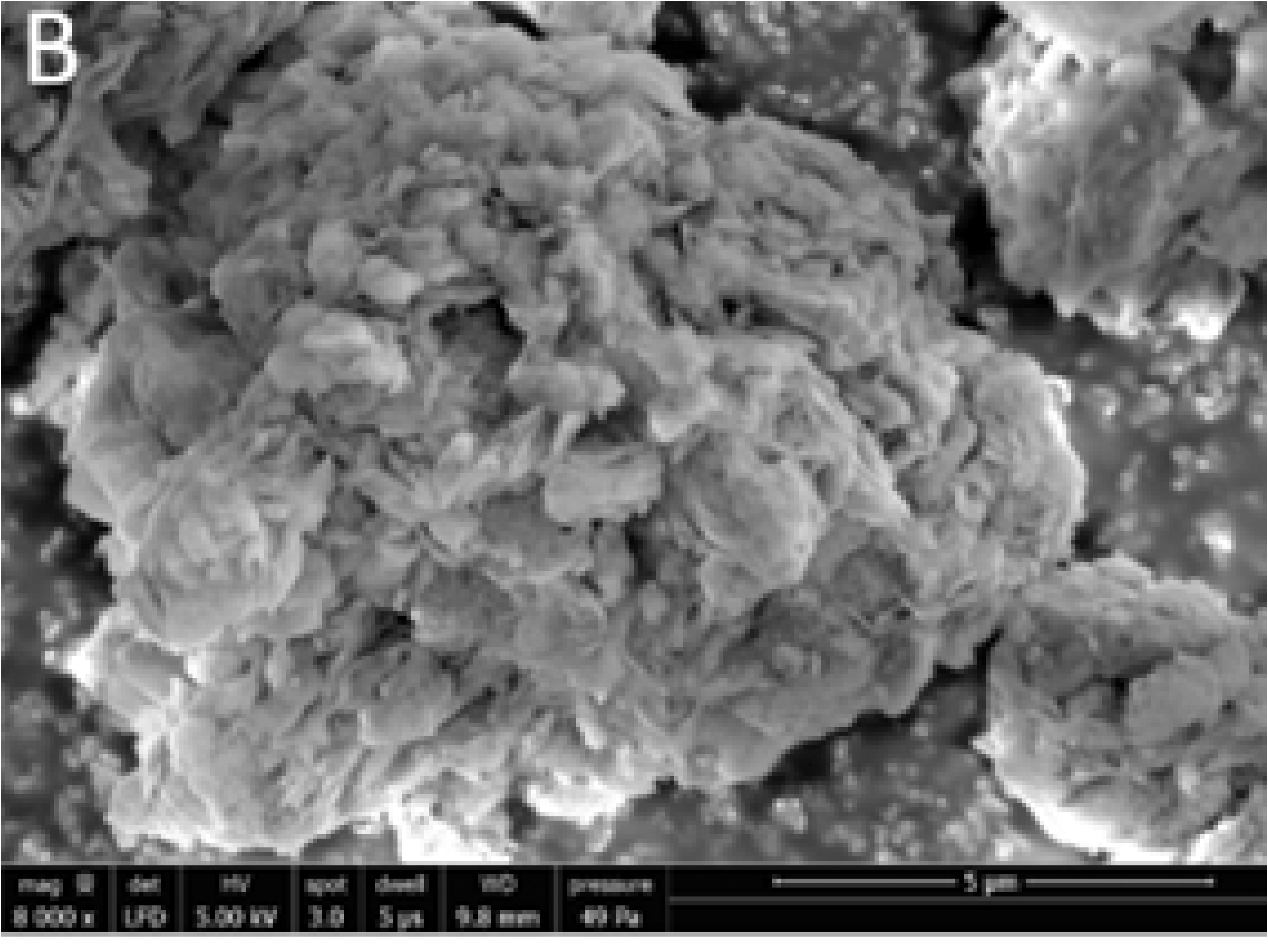
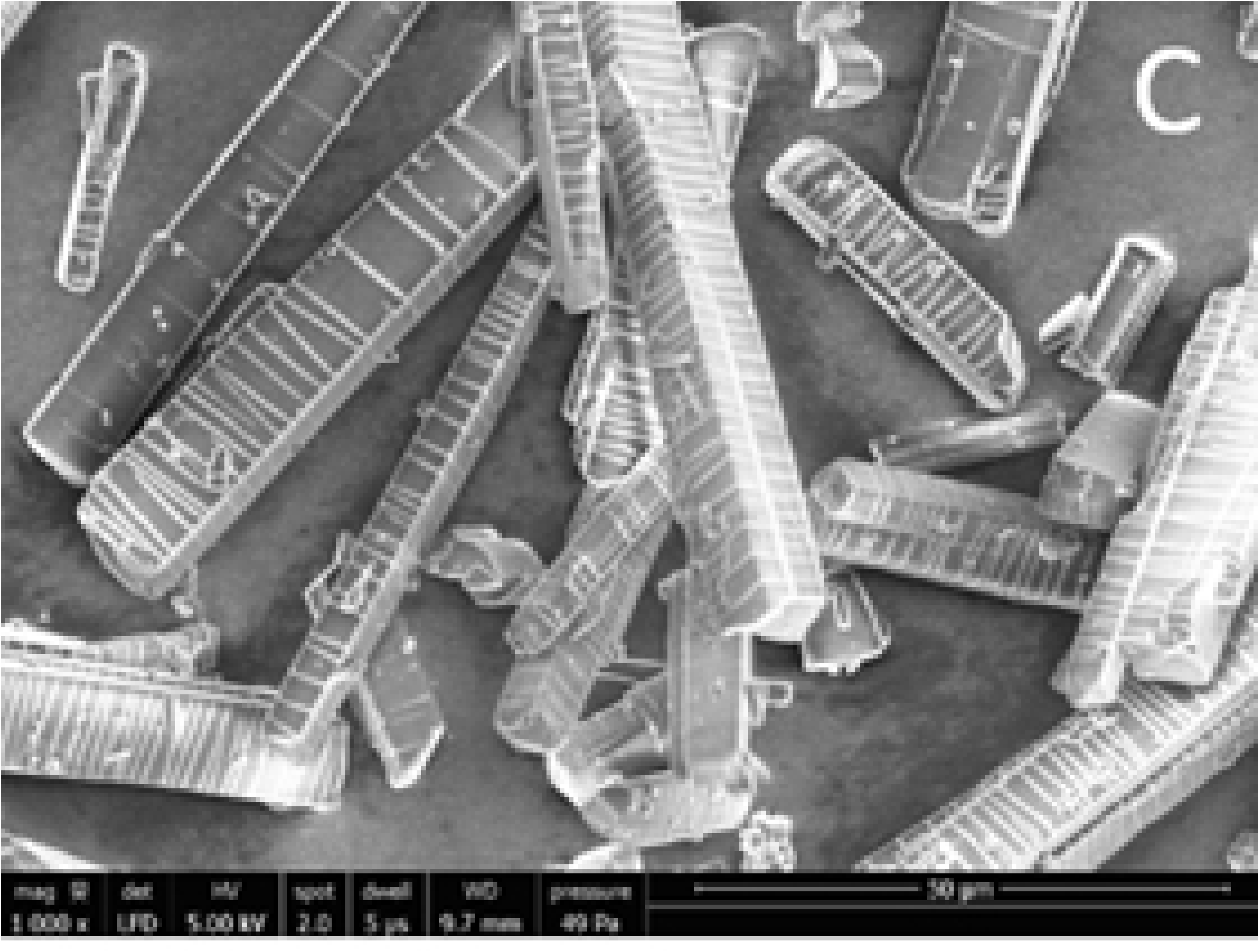
SEM images of a) FCC fines, b) 15 w/w% amoxicillin-loaded FCC fines, and c) powder amoxicillin.

This morphological difference suggests that the amoxicillin, when loaded into the FCC fines during the solvent evaporation process, precipitates in an amorphous or 2D nanocrystallite-composite sheet state. This altered structural state is likely a key factor contributing to the observed enhanced dissolution rate of the drug. We can speculate that the driving mechanism for nanocrystal formation could be related to a sorption effect of molecules in contact with a surface which could provide a nanoscale molecular orientation effect, a conformation that prevents the lowest energy intermolecular structure, namely preventing crystal formation.

### 3.4. Release of amoxicillin from the EC films in water

From the wide range of polymers suitable for a polymer matrix, we selected ethyl cellulose (EC), a non-biodegradable, biocompatible polymer that is one of the most studied encapsulating materials for controlled drug release (9, 22–25). Although cellulose and its derivatives are environmentally friendly and actively degradable by various bacteria and fungi, Marston *et al*. demonstrated that cellulose is a biodurable material when implanted in animal and human tissues, as resorption does not occur due to the absence of cellulase synthesis in cells (26–28). Please note that for these experiments we used the Omyapharm particles – which are the starting material for the FCC fines and granules.

Depending on intended application, the amoxicillin release can be adjusted in various ways, such as: by tuning the amoxicillin loading level in the porous particles, changing the amount of particles in the EC solution formulation, as well as by inclusion of additives. In this proof of concept experiments we show the release of amoxicillin at a specific surface to water exposure.

In this case a rolled EC film was immersed in water where the surface to water exposure was set to 5.4 cm^2^/mL, as described in the Materials section. To evaluate the advantage of loading amoxicillin in porous particles and the influence of an additional loading step, the EC film was compared to a reference EC film containing a physical mixture of amoxicillin and Omyapharm particles where the amoxicillin and Omyapharm were added separately into the polymer solution, as seen in Figure 8. Note that the ratio of all components was kept identical.

**Figure 8.**
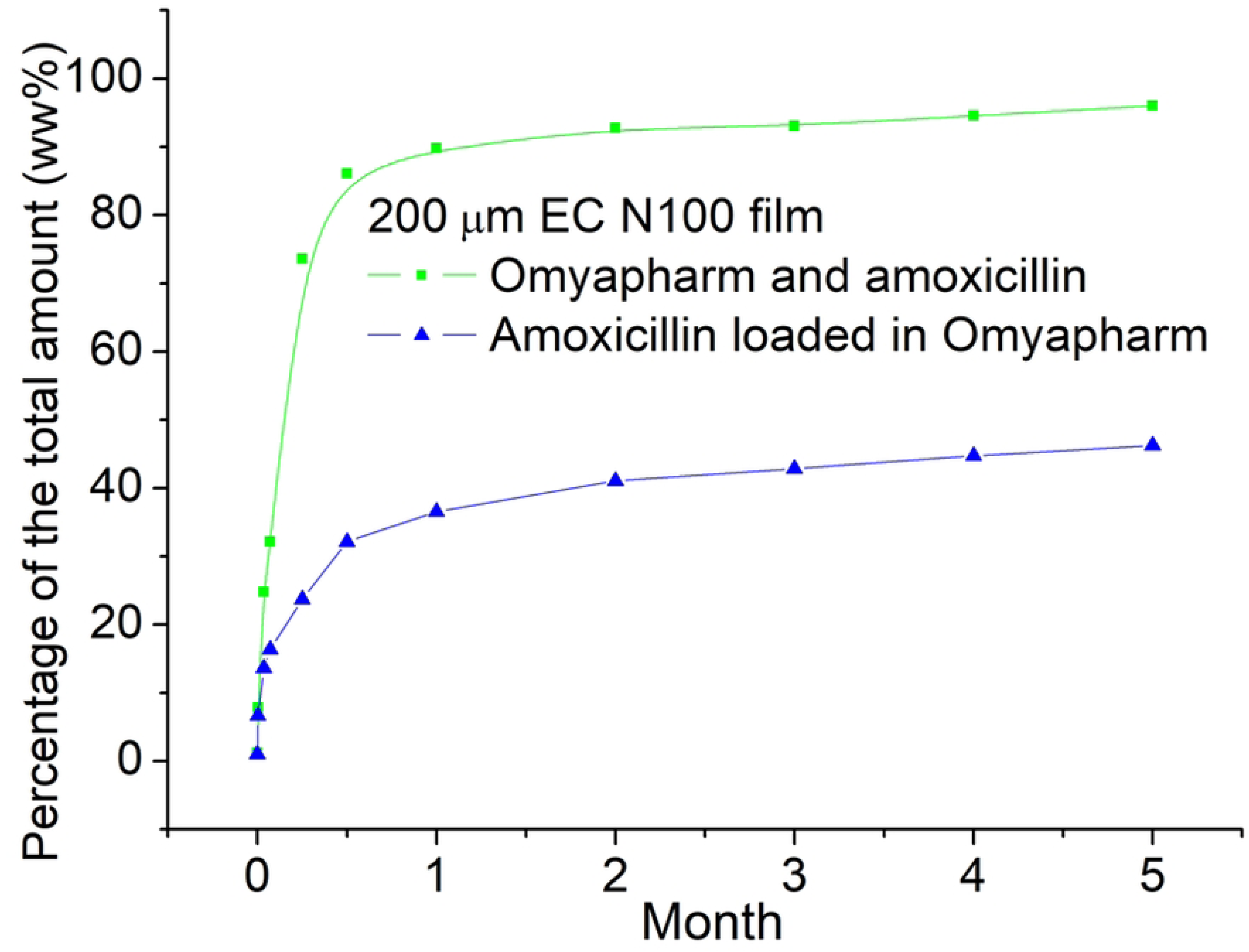
Amoxicillin release over time from the following films: Omyapharm particles loaded with amoxicillin (blue), Omyapharm particles and amoxicillin mixture (green) - as a reference.

In the release experiments, the amoxicillin concentrations were measured after 2 h, 1 day and 2 days, 1 week, 2 weeks 4 weeks and then monthly up to 5 months. As sown in Figure 9, both release profiles show burst behavior over the first approximative 15 days reaching 85% for unloaded amoxicillin (physical mixture) and 39% for loaded amoxicillin, respectively. After approximatively 1 month, both profiles show similar behavior with a slight monotonous increase kinetics. At the end of the experiment (5 months), unloaded amoxicillin was almost completely released (96%) whereas films with loaded amoxicillin had only released 46% of the amoxicillin.

**Figure 9.**
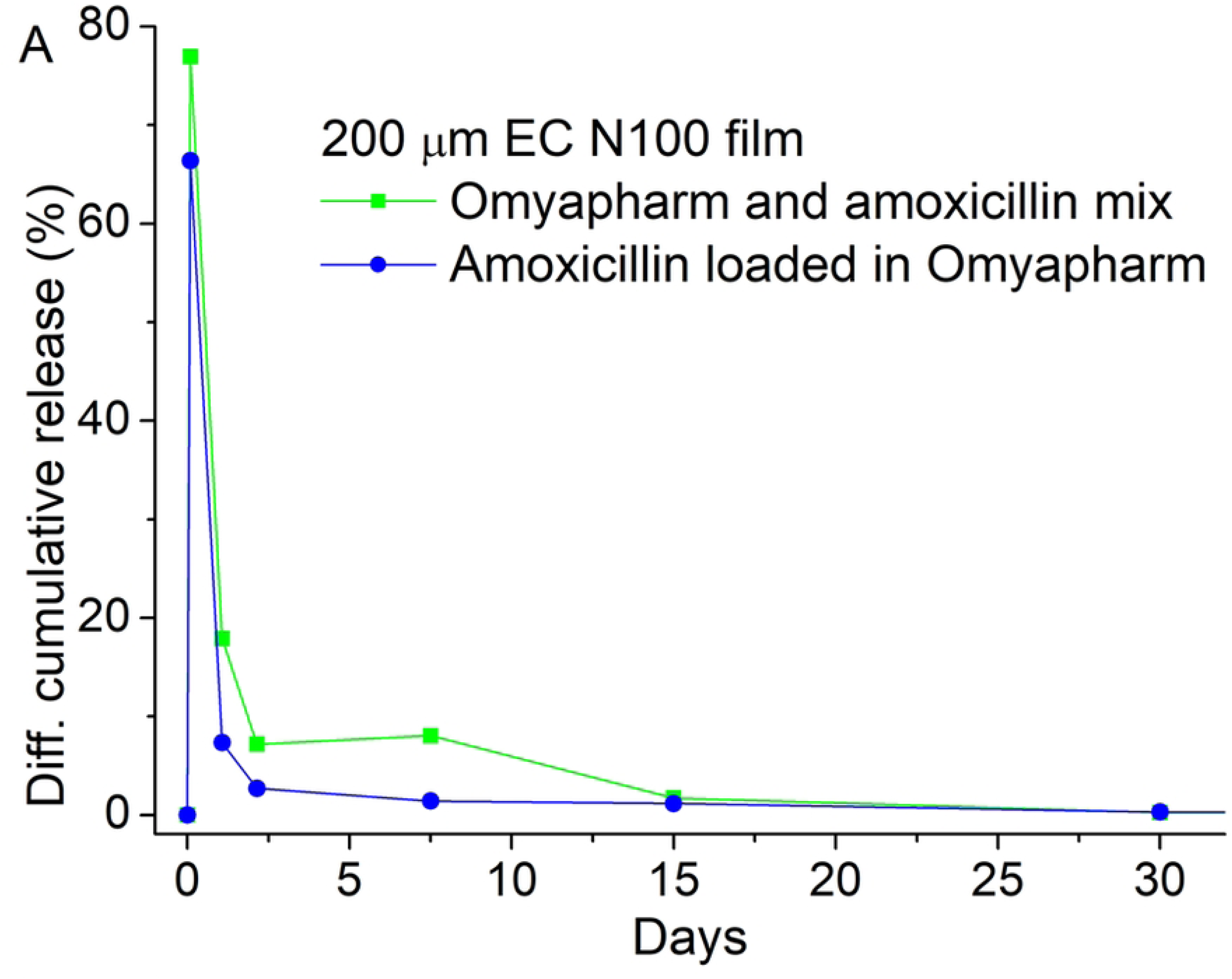
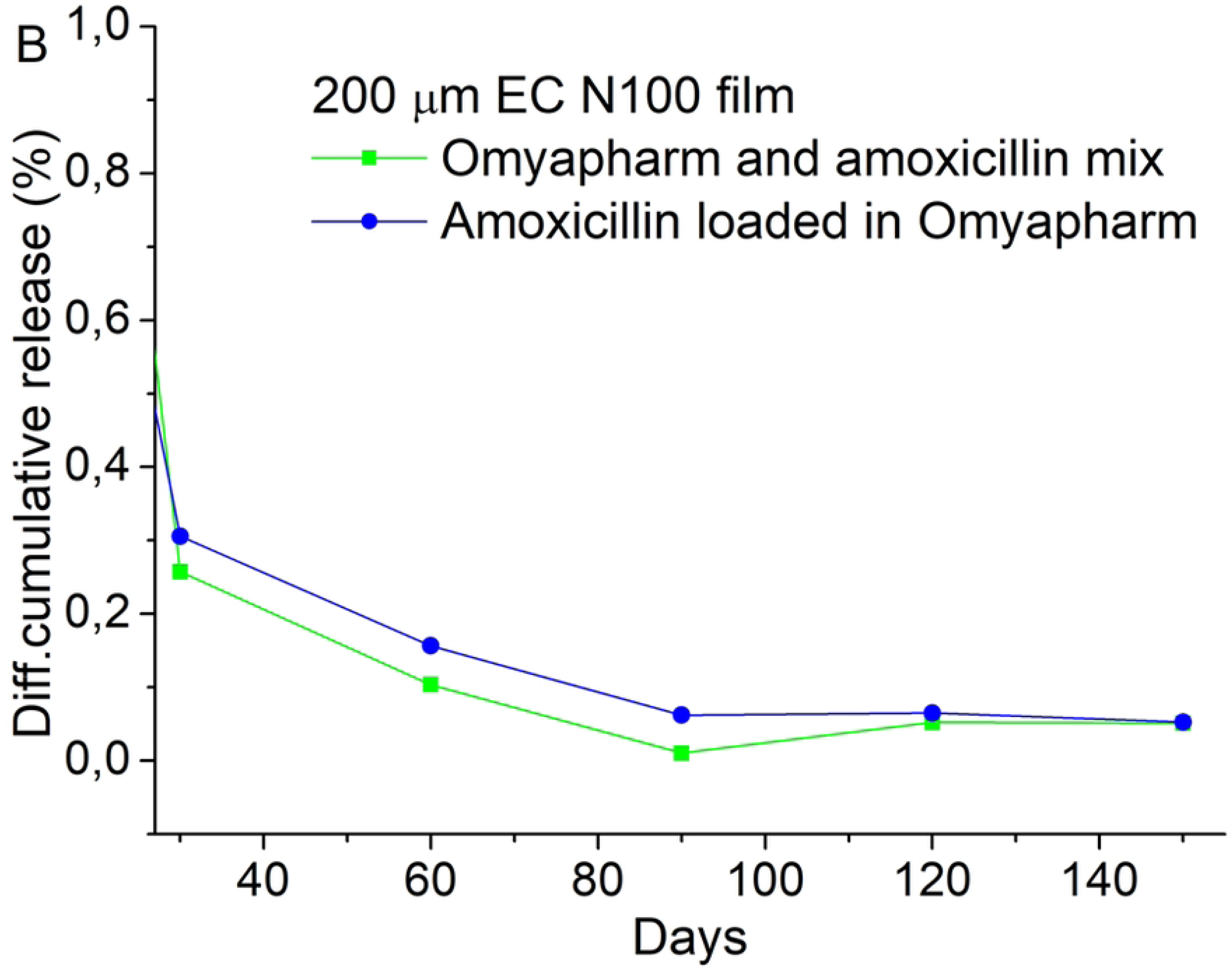
The differential release profile of the films made from the amoxicillin loaded in Omyapharm particles (blue) and Omyapharm particles and amoxicillin mixture (green) for A) month 1 and B) month 2-5.

By closely looking at the release rate, it can be noted that in the second phase of the release (month 2 to 5) amoxicillin loaded into Omyapharm is released at a higher rate than the unloaded part. During the initial burst phase (month 1), the average release rate is 13 %/month for the loaded amoxicillin and 19 %/month for the unloaded amoxicillin, respectively. During the second phase (month 2 to 5), the average release rate is 0.032 %/month for the loaded amoxicillin and 0.023 %/month for the unloaded amoxicillin, respectively. In other words, the extended release appears to be a result of a reduced burst rather than a slower diffusion of amoxicillin through the EC film. Loading actives into porous Omyapharm particles prior to the integration into a polymer matrix might therefore be a viable approach to mitigate extensive burst of actives from thin films.

Albeit a different dosage form, this time frame is also much longer compared to the Singh *et al.* study (5), where release of amoxicillin from microspheres was sustained by/through a mucoadhesive device. Their best formulation, based on a combination of HPMC K4M as mucoadhesive and Eudragit RS 100 as polymer matrix, showed a continuous release over a maximum of 12 h only in-vitro, with the release profile reaching 96.15 w/w% of the total amount in that relatively short period. Also Torres-Figueroa *et al.* (6) have studied the effectiveness of composite hydrogels - poly(acrylamide) and starch - as a platform for controlled release of amoxicillin, however the hydrogel reported released 48.6 w/w% of the amoxicillin load within the first 24 h. This behavior was attributed to the hydrophilic character of the starch and the swelling rate of the hydrogel affecting the kinetics of the drug release. The longest sustained drug release we found in the literature was obtained by Prasanna *et al.* (8), who studied the release of amoxicillin from a hydroxyapatite layer-by-layer coated with polyvinyl alcohol and sodium alginate. Apart from a good antibacterial activity they showed a sustained release of amoxicillin for about 30 days. In none of these studies did the loading/impregnation of amoxicillin change the physical state from the original crystalline form, in contrast to where we see the loading of Omyapharm particles resulting in a change from long range crystalline to a pseudo amorphous state.

The amoxicillin release depends on the hydrophilic character of the used polymer. HPMC (5), poly(acrylamide), starch (6) or polyvinyl alcohol (8) have been reported examples. In our case, although EC is not water soluble, it is nonetheless hydrophilic. In spite of the insolubility of EC, the amoxicillin finds its way out to the surface along hydrophilic fissures, most likely formed during the initial step of solvent evaporation that takes place during film formation. Relying on such fissures would not be a guarantee for successful delivery.

Another more likely possibility is that water molecules, under osmotic pressure, can diffuse into the molecular-sized spaces of EC polymer and reach amoxicillin loaded particles. Thus, the amoxicillin solution formed within the film can diffuse out under the resulting concentration gradient. Even though the amoxicillin loaded in the porous particles have a higher water solubility (dissolution rate) as compared to pure amoxicillin, the results of the tempered release rate of the loaded amoxicillin can be mainly explained by the significantly reduced initial burst. However, additional contribution may come from other factors such as: i) additional tortuosity provided by the particle; ii) sedimentation of the loaded particles (higher density provided by the porous particles) during film production and hence longer diffusion pathway through the EC matrix; iii) easier access to the more homogeneously distributed and smaller pure amoxicillin particles as compared to more concentrated amoxicillin – porous particles loaded domains, which, in turn, can be more effectively encapsulated by the EC.

The results obtained in this study suggest that a more efficient system in terms of controlled delivery can be obtained by loading amoxicillin into porous microparticles – functionalized calcium carbonate – instead of mixing it directly into a polymer matrix; and then homogeneously distributing these particles within a polymer matrix. Reducing initial burst avoids unnecessary high concentrations and allows for more judicious release rates over a prolonged period of time. The observed release concentration of amoxicillin (above 0.1 mg/mL) exceeds the minimum inhibitory concentration (MIC) for E. coli, which is 0.008 mg/mL (29, 30). Eventually, the release rates of amoxicillin from ethyl cellulose films can be tailored by manipulating the preparation method, film thickness, and the inclusion of specific additives in order to achieve relevant performance as drug delivery systems in catheter and indwelling applications.

## 4. Conclusions

Since bacterial resistance to antibiotics is recognized currently as a major threat to global health it is desirable to change the way antibiotics are used, crucially supporting their judicious application. Today, already established infections can only be treated with selected antibiotics, and it is important to use as little as possible to minimize antimicrobial resistance, but the key point is that this controlled small amount must be maintained for long periods above the minimum inhibition concentration (MIC). Therefore, sustained release systems are regarded as a viable solution to maintaining microbial-free treatment and, hence, reducing the chance of antimicrobial resistance.

Within this work, a model drug – amoxicillin, was successfully loaded into porous functionalized calcium carbonate particles using a solvent evaporation method achieving drug loads of 30 wt%. Dissolution of loaded amoxicillin was faster compared to pure amoxicillin as demonstrated by UV-vis spectrometry. This can be explained by the inhibition of crystal formation during the loading process as supported by XRD and SEM measurements. Structural analysis reveals the presence of a pseudo amorphous, nanocrystalline or molecularly surface oriented mesophase layer form. However, a total area of only 2880 µm^2^ was investigated by nanoscale XRD within this study, therefore the reported observations are only qualitative interpretations. Further measurements of larger areas and on more samples could produce more statistics, thus leading to quantitative results, such as the relative abundance of nanocrystalline or mesophase areas with respect to amorphous ones as well as the average size of nanocrystals or mesophase layers.

If this essentially non-crystalline material is embedded in a biocompatible polymer, such as ethyl cellulose (EC), a flexible water insoluble film, that can be easily manipulated, could be produced. With this film construct we have shown that the amoxicillin release can be sustained for more than 5 months. The release through the EC matrix of amoxicillin from loaded particles may effectively mitigate the issue of burst release by minimizing the initial surge and ensures a sustained release of the active ingredient over an extended period, reducing wastage during the early phase. In other words, the same amount of amoxicillin can be delivered over a much longer period of time, rendering this film well suited, for example, in catheter/indwelling applications.

## 5. ACKNOWLEDGEMENT

This work was supported by Omya International AG and RISE Research Institute of Sweden. Ulf Johansson is gratefully acknowledged for support during the XRD experiment at NanoMAX.

## Notes

### Competing Interest Statement

The authors have declared no competing interest.

## References

1. van der Horst MA, Schuurmans JM, Smid MC, Koenders BB, ter Kuile BH. De novo acquisition of resistance to three antibiotics by Escherichia coli. Microb Drug Resist. 2011;17(2):141–7.

2. Gao P, Nie X, Zou M, Shi Y, Cheng G. Recent advances in materials for extended-release antibiotic delivery system. The Journal of Antibiotics. 2011;64(9):625–34.

3. Smith FPDJHCNH, inventor; Beecham Research Laboratories Ltd, assignee. Improvements in or relating to penicillin derivatives. UK1958.

4. Akhavan BJ KN, Vijhani P. Amoxicillin in StatPearls. Treasure Island (FL). https://www.ncbi.nlm.nih.gov/books/NBK482250/: StatPearls Publishing; 2021.

5. Singh S, Chidrawar V, Ushir Y, Vadalia K, Sheth N, Shukla SHN. Pharmaceutical Characterization of Amoxicillin Trihydrate as Mucoadhesive Microspheres in Management of H. Pylori. International Journal of PharmTech Research; 2003. p. 348–58.

6. Ana V. Torres-Figueroa CJP-M, Teresa del Castillo-Castro, Enrique Bolado-Martínez, María A. G. Corella-Madueño, Alejandro M. García-Alegría, Tania E. Lara-Ceniceros, Lorena Armenta-Villegas,. Composite Hydrogel of Poly(acrylamide) and Starch as Potential System for Controlled Release of Amoxicillin and Inhibition of Bacterial Growth. Journal of Chemistry. 2020;2020:14.

7. Songsurang K, Pakdeebumrung J, Praphairaksit N, Muangsin N. Sustained release of amoxicillin from ethyl cellulose-coated amoxicillin/chitosan-cyclodextrin-based tablets. AAPS PharmSciTech. 2011;12(1):35–45.

8. Prasanna APS, Venkatasubbu GD. Sustained release of amoxicillin from hydroxyapatite nanocomposite for bone infections. Progress in Biomaterials. 2018;7(4):289–96.

9. Andersen MJ, Flores-Mireles AL. Urinary Catheter Coating Modifications: The Race against Catheter-Associated Infections. Coatings. 2020;10(1):23.

10. Merchant J, Müllertz A, Rades T, Bannow J. Functionalized calcium carbonate (FCC) as a novel carrier to solidify supersaturated self-nanoemulsifying drug delivery systems (super-SNEDDS). European Journal of Pharmaceutics and Biopharmaceutics. 2023;193:198–207.

11. Lundin Johnson M, Noreland D, Gane P, Schoelkopf J, Ridgway C, Millqvist Fureby A. Porous calcium carbonate as a carrier material to increase the dissolution rate of poorly soluble flavouring compounds. Food & Function. 2017;8(4):1627–40.

12. Levy CL. Nanoporous Calcium Carbonate-Based Substrates for the Controlled Delivery of Functional Materials. Plymouth: University of Plymouth; 2017.

13. Johansson U, Carbone D, Kalbfleisch S, Bjorling A, Kahnt M, Sala S, et al. NanoMAX: the hard X-ray nanoprobe beamline at the MAX IV Laboratory. Journal of Synchrotron Radiation. 2021;28(6):1935–47.

14. Robert A, Cerenius Y, Tavares PF, Hultin Stigenberg A, Karis O, Lloyd Whelan A-C, et al. MAX IV Laboratory. The European Physical Journal Plus. 2023;138(6):495.

15. Björklund OSE. Thermal stability assessment of antibiotics in moderate temperature and subcritical water using a pressurized dynamic flow-through system. International Journal of Innovation and Applied Studies. 2015;11(4):872–80.

16. Boles MO, Girven RJ, Gane PAC. The structure of amoxycillin trihydrate and a comparison with the structures of ampicillin. Acta Crystallographica Section B. 1978;34(2):461–6.

17. Bird AE. Amoxicillin. In: Brittain HG, editor. Analytical Profiles of Drug Substances and Excipients. 23: Academic Press; 1994. p. 1–52.

18. U.S. PC. Product Information Report: Amoxicillin. Rockville, Maryland: Pharmacopeial Convention U.S.; 2017.

19. Rowe MC, Brewer BJ. AMORPH: A statistical program for characterizing amorphous materials by X-ray diffraction. Computers & Geosciences. 2018;120:21–31.

20. Chung FH. Unified Theory for Decoding the Signals from X-Ray Florescence and X-Ray Diffraction of Mixtures. Applied Spectroscopy. 2017;71(5):1060–8.

21. Jenkins R. X-Ray Techniques: Overview. Encyclopedia of Analytical Chemistry.

22. Sánchez-Lafuente C, Teresa Faucci M, Fernández-Arévalo M, Álvarez-Fuentes J, Rabasco AM, Mura P. Development of sustained release matrix tablets of didanosine containing methacrylic and ethylcellulose polymers. International Journal of Pharmaceutics. 2002;234(1):213–21.

23. Wasilewska K, Winnicka K. Ethylcellulose-A Pharmaceutical Excipient with Multidirectional Application in Drug Dosage Forms Development. Materials (Basel). 2019;12(20):3386.

24. Notario-Pérez F, Cazorla-Luna R, Martín-Illana A, Galante J, Ruiz-Caro R, das Neves J, et al. Design, fabrication and characterisation of drug-loaded vaginal films: State-of-the-art. Journal of Controlled Release. 2020;327:477–99.

25. Murtaza G. Ethylcellulose microparticles: a review. Acta Pol Pharm. 2012;69(1):11–22.

26. Sannino A, Demitri C, Madaghiele M. Biodegradable Cellulose-based Hydrogels: Design and Applications. Materials. 2009;2(2):353–73.

27. Elçin AE. In Vitro and In Vivo Degradation of Oxidized Acetyl- and Ethyl-Cellulose Sponges. Artificial Cells, Blood Substitutes, and Biotechnology. 2006;34(4):407–18.

28. Märtson M, Viljanto J, Hurme T, Laippala P, Saukko P. Is cellulose sponge degradable or stable as implantation material? An in vivo subcutaneous study in the rat. Biomaterials. 1999;20(21):1989–95.

29. Stohr JJJM, Kluytmans-van den Bergh MFQ, Verhulst CJMM, Rossen JWA, Kluytmans JAJW. Development of amoxicillin resistance in Escherichia coli after exposure to remnants of a non-related phagemid-containing E. coli: an exploratory study. Antimicrob Resist Infect Control. 2020;9(1):48-.

30. EUCAST ECoAST-. Antimicrobial wild type distributions of microorganisms https://mic.eucast.org/search/:eucast.org; 2021 [Web Database].

